# Heterogeneity in the response of different subtypes of *Drosophila melanogaster* enteroendocrine cells to viral infections

**DOI:** 10.1101/2020.07.01.182352

**Authors:** João M. F. Silva, Tatsuya Nagata, Fernando L. Melo, Santiago F. Elena

## Abstract

Single cell RNA sequencing (scRNA-seq) offers the possibility to monitor both host and pathogens transcriptomes at the cellular level. Here, public scRNA-seq data from *Drosophila melanogaster* have been used to compare the differences in replication strategy and cellular response between two viruses, Thika virus (TV) and *D. melanogaster* Nora virus (DMelNV) in enteroendocrine cells (EEs). TV and DMelNV exhibited different patterns of replication and for TV, accumulation varied according to cell subtype. Cells infected with TV underwent down-regulation of genes that represent bottlenecks in the fruit fly interactome, while cells infected with DMelNV went through a down-expression of translation-related genes that represent both hubs and bottlenecks in the host interactome. In contrast, flies infected with DMelNV show only a systemic level down-regulation of bottleneck genes. Here, we use scRNA-seq to highlight the differences and commonalities between cellular response to TV and DMelNV and between cellular and systemic response to DMelNV.

## Introduction

Enteroendocrine cells (EEs) play an important role in the control of many physiological processes. They are scattered throughout the gastrointestinal tract, making it the largest endocrine organ in the body, and produce more than 20 neuropeptide hormones including allatostatines A, B and C (AstA, AstB and AstC, respectively), tachykinin (Tk), neuropeptide F (NPF), diuretic hormone 31 (DH31), and CCHamide-1 and −2 (CCHa1 and CCHa2, respectively) (Rehfeld, 2004; Furness et al., 2013; Gribble and Reimann, 2019; Beehler-Evans and Micchelli, 2015). Recently, ten EE subtypes were identified in the fruit fly *Drosophila melanogaster* Meigen. These subtypes can be divided in two major classes: class I is composed by cells expressing AstC, and class II is composed by cells that express Tk (Beehler-Evans and Micchelli, 2015; Guo et al., 2019). In addition to these classes, a subtype dubbed III that does not express neither AstC nor Tk was also identified. Further classification of EE subtypes is based on whether they are located in the anterior, medium (gastric region) or posterior regions of the gastrointestinal tract (-a, -m and -p suffixes; Figure 1A) and on their gene expression (Guo et al., 2019).

**Figure 1.**
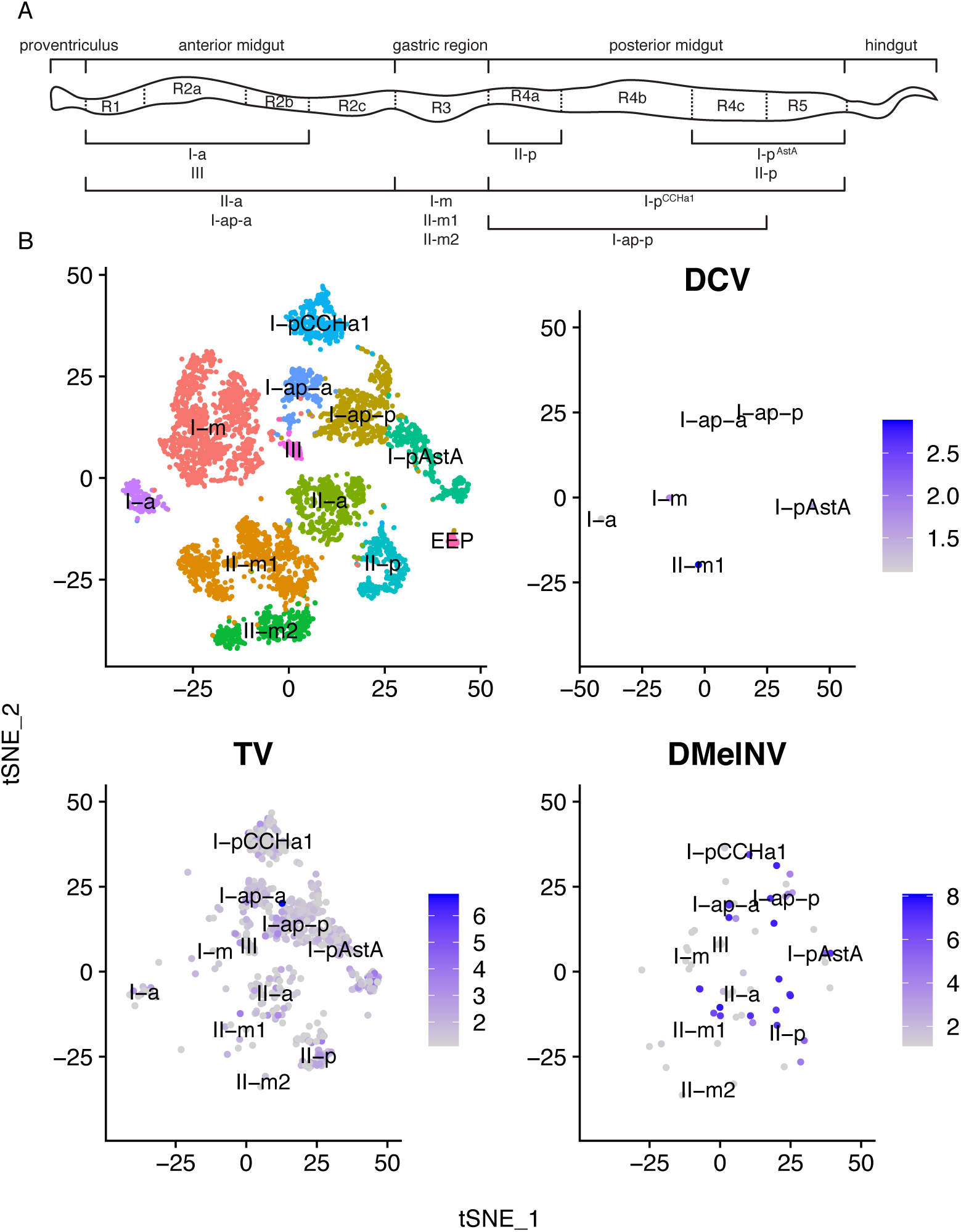
Detection of RNA viruses in a public EE *D. melanogaster* scRNA-seq dataset. (A) Schematic representation of the *D. melanogaster* gastrointestinal tract showing the spatial distribution of EEs (adapted from Guo et al., 2019). (B) t-SNE (*t*-distributed stochastic neighbor embedding) reduction plots of EEs with identified clusters; and cells infected with DCV, TV and DMelNV, along with their log-expression (accumulation) level.

The fruit fly is an attractive invertebrate model for studying virus-host interactions (Huszar and Imler, 2008), in which RNA interference (RNAi) plays a major antiviral role (Kemp and Imler, 2009; Van Mierlo et al., 2012). Novel sequencing technologies are allowing the discovery of novel RNA viruses in both wild and stock flies (Webster et al., 2015) that may serve as models for studying virus-host interactions. However, the biology of only a small subset of these viruses has been investigated in depth, such as *D. melanogaster* sigma virus (DMelSV; genus *Sigmavirus*, family *Rhabdoviridae*), drosophila C virus (DCV; genus *Cripavirus*, family *Dicistroviridae*), and more recently, *D. melanogaster* Nora virus (DMelNV; unclassified picorna-like virus), an enteric virus that is known to cause persistent infections on both wild and stock flies with no obvious pathological outcome (Huszar and Imler, 2008; Habayeb et al., 2006; Habayeb et al., 2009; Webster et al., 2015).

Single-cell RNA-sequencing (scRNA-seq) allows for the analysis of subtle and cell (sub)type-dependent alterations of gene expression that would otherwise be undetected in bulk RNA-seq experiments. Here, cells infected with picorna-like viruses from a publicly available fly EEs scRNA-seq dataset (Guo et al., 2019) were used to investigate the infection dynamics of two viruses, DMelNV and the recently discovered Thika virus (TV; unclassified picorna-like virus), on different subtypes of EEs. Since these viruses are polyadenylated, we have been able of quantifying their replication level based on their number of unique molecular identifiers (UMI). We found a drastic contrast in the replication pattern between these two viruses. On the one hand, DMelNV only infected a few cells but exhibited high expression levels that sometimes exceeded 50% of the total mRNA of the cell while, on the other hand, TV was able to infect a higher number of cells, but its replication level was generally low. Cells infected with either DMelNV or TV exhibited both generic and cell-subtype-specific transcriptional responses, and most important, they responded differentially to DMelNV or TV infection. DMelNV infection led to a down-regulation of essential hubs and bottlenecks of interactions on the *D. melanogaster* interactome, which are mostly composed by ribosomal proteins. By analyzing a publicly available bulk RNA-seq data from DMelNV-infected flies (Lopez et al., 2018), we found that in this systemic response to DMelNV, only bottleneck genes were down-regulated, and no significant perturbation of hubs was found. Similarly, cellular response to TV infection was characterized by a down-regulation of bottleneck genes only.

By analyzing both viral replication and host transcription levels, in a manner that is analogous to dual RNA-seq (Wesolowska-Andersen et al., 2017), we show that in the presence of multiple infections it is possible to use scRNA-seq to analyze the transcription of the host and multiple pathogens. Here, not only we have been able to monitor both virus replication and host transcriptional levels, we have been also able to simultaneously compare the replication and virus-host interaction strategies of two different viruses *in vivo*, revealing more of the uniqueness and similarities of viral infections at the single-cell level and providing possible new models for invertebrate viruses.

## Results

### Detection of RNA viruses in public *D. melanogaster* scRNA-seq datasets

A publicly available EE scRNA-seq dataset from *D. melanogaster* (BioProject accession: PRJNA547484) was obtained from the Sequence Read Archive (SRA; https://www.ncbi.nlm.nih.gov/sra) and investigated for the presence of viruses with BLASTn (Altschul et al., 1990). Three RNA viruses were identified: DCV, DMelNV and TV. Next, reads were aligned to the *D. melanogaster* and virus genomes to obtain gene/viral counts for each cell, which were used to cluster generation. All clusters were identified based on a set of marker genes (Guo et al., 2019; Figure S1A). In total, six, 67 and 794 cells infected with DCV, DMelNV, and TV, respectively, were found. Figure 1A shows a schematic representation of *D. melanogaster* gastrointestinal tract, along with the spatial distribution of EEs subtypes. Figure 1B shows t-SNE reduction plots of the cell clusters, together with the log-expression values of DCV, DMelNV and TV. No infected EE progenitor (EEP) could be found, and no cell from the I-a subtype was found to be infected with DMelNV. We also found 13 cells coinfected with TV and DMelNV. Given the low incidence of DCV infections, we discarded this virus from subsequent analyses.

In addition to the EE scRNA-seq dataset, a publicly available scRNA-seq dataset from the entire midgut of fruit flies was also investigated for the presence of viruses (BioProject accession: PRJNA493298). In this data, to which we will refer as midgut atlas dataset, only DMelNV was detected. Like the EE dataset, cell clusters were generated and identified based on a set of marker genes (Hung et al., 2020; Figure S1B). Major well defined clusters composed by enterocytes (ECs) from the anterior, middle and posterior portions of the midgut (aECs, mECs and pECs, respectively) were identified, along with clusters composed by cardia cells, located in the cardia (proventriculus) region; intestinal stem cells and enteroblasts (ISC/EBs); large flat cells (LFCs), which are located in the posterior gastric region; copper/iron cells, located in the gastric region; and one cluster composed by unknown cell types (unk) (Figure S2A). Clusters composed by differentiating ECs and EC-like cells (Hung et al., 2020) were not identified, possibly due to removal of these cells in our pre-processing step and/or due to them clustering to their most similar ECs. A higher prevalence of DMelNV was noted in this dataset, with 994 cells infected with this virus. Throughout this work the midgut atlas dataset was used to complement and confirm the results obtained for DMelNV on the EE dataset.

### DMelNV and TV exhibit different patterns of replication

The replication level of both DMelNV and TV was evaluated by analyzing both the percentage of viral RNA in the cells and log-normalized expression values (Figure 2A and B). A dramatic contrast in the replication levels between the two viruses was found. TV infected a higher number of cells compared to DMelNV, but exhibited lower replication levels per cell. The percentage of viral RNA in TV infected cells was always below 5% with only one exception (a single cell I-ap-p), whereas for DMelNV, the percentage of viral RNA was up to ∼80%. The percentage of infected cells on each cluster also varied drastically between these viruses (Figure 2C). TV infected a very low proportion of cells from subtypes located in the gastric region, and was found on more than 60% of the cells of the I-p^AstA^ cell-subtype located in the posterior region of the gastrointestinal tract that is characterized by the expression of AstC, AstA and CCHa1.

**Figure 2.**
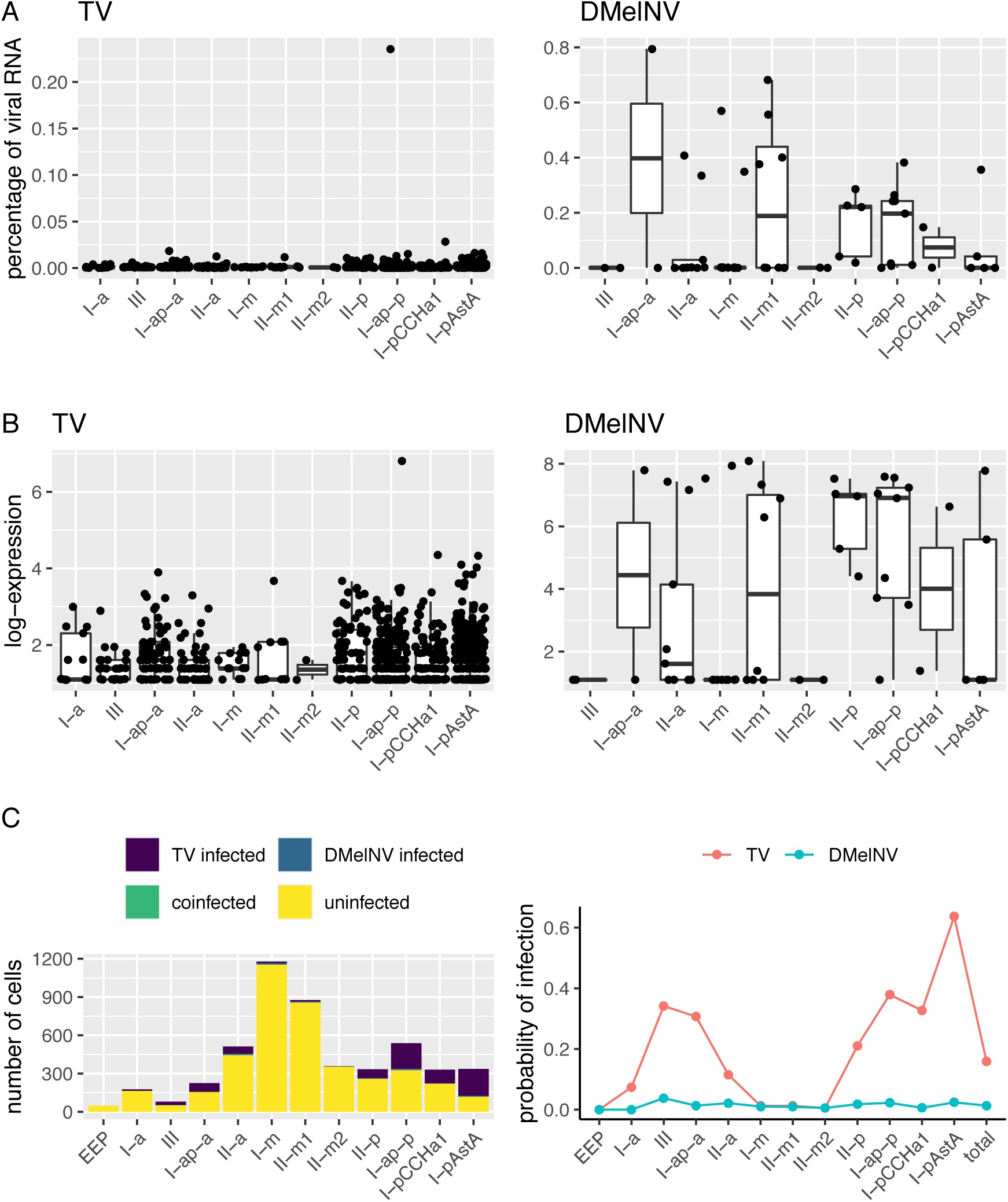
TV and DMelNV exhibit different patterns of replication. (A) Boxplots showing the percentage of viral RNA from TV and DMelNV for each EE subtype. Each point represents an individual cell. (B) Boxplots showing log-expression values of TV and DMelNV for each EE subtype. Each point represents an individual cell. (C) Number (left) and percentage (right) of cells from each subtype infected with TV and DMelNV.

One-way ANOVA tests showed a significant influence of EE subtype on the expression level of TV (*P* = 7.3×10^−4^) but not of DMelNV (*P* = 0.12). Following these results, Tukey’s HSD *post hoc* tests revealed two well defined groups exhibiting, respectively, the highest and lowest levels of replication for TV: one composed by EE subtypes II-p and I-p^AstA^, both located in the posterior part of the gastrointestinal tract; and other composed by EE subtypes II-m, located in the gastric region, and II-a and III, both located in the anterior region of the gastrointestinal tract (Figure 1A and 2C). In accordance with the results obtained for the EE dataset, no difference in replication level according to cell subtype was detected for DMelNV when analyzing the midgut atlas dataset (one-way ANOVA; *P* = 0.25).

### Expression of translation-related genes is negatively correlated to DMelNV replication level

We found 88 and four genes whose expression levels were significantly correlated to the replication level of DMelNV and TV, respectively. Possibly, the narrow range of variation of the replication level of TV hampered the detection of host correlated genes. The expression of one gene, a mitochondrial large RNA (mt:lrRNA), was positively correlated to the replication of TV (adjusted *P* = 9.7×10^−7^), whereas a negative correlation was found for tetraspanin 42Ec (Tsp42Ec; *P* = 0.014), CDC50 (*P* = 0.022) and CG14989 (*P* = 0.004). The most significant correlation was found for mt:lrRNA, which plays a role in the formation of granular poles in germ line progenitor cells (Iida and Kobayashi, 1998). GO enrichment analyses were performed for nine and 79 genes which were positively and negatively correlated, respectively, to the replication level of DMelNV (Figure 3). An overrepresentation in processes mostly related to translation was found in negatively correlated genes but no overrepresentation was found for the few positively correlated genes. For the midgut atlas dataset, the expression of 112 genes was found to be correlated to the replication level of DMelNV, with 27 correlated genes found in both datasets (supplementary file S1). Similar GO terms were obtained for negatively correlated genes for the midgut atlas dataset, although processes related to response to inorganic response, import into cell and others were enriched for positively correlated genes (Figure S3). Given the high number of DMelNV-infected cells and the high variability in the replication levels of DMelNV in this dataset, the results obtained from the correlation analysis may be a powerful mean to uncover generic cellular response to this virus.

**Figure 3.**
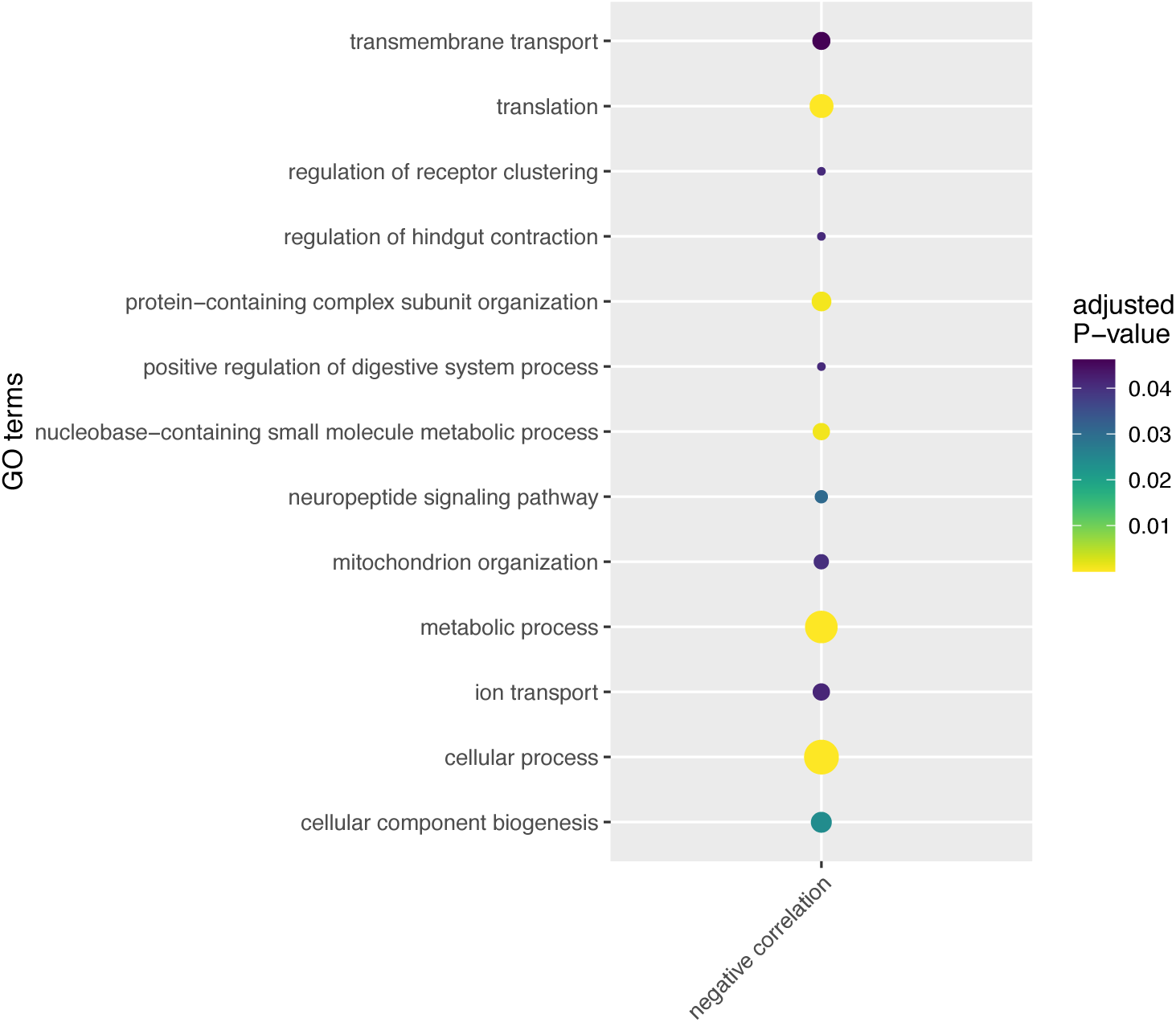
DMelNV replication correlation to gene expression. GO enrichment analysis of genes which expression is negatively correlated to the replication level of DMelNV for the EEs dataset. The size of the points represents the number of times the respective term was present.

### Generic and cell-subtype-specific transcriptional response to TV and DMelNV

In addition to the gene expression correlation analyses we also investigated differentially expressed genes (DEGs) according to cell infection status (infected or uninfected) and the interaction between infection status and cell-subtype. In the first class, which we will refer to as generic response genes, we found 126 and 32 DEGs for DMelNV and TV, respectively; whereas in the second class, referred to as cell-subtype-specific response genes, we found 21 and 13 DEGs for DMelNV and TV, respectively. These categories are not exclusive, and some DEGs are also correlated with the replication level of the virus (supplementary file S1). Inspection of the cell-subtype-specific response genes revealed that, in most cases, cell-subtypes from the gastric region responded differently to TV (Figure S4A). A similar pattern was not obvious for DMelNV (Figure S4B). In a few cases, like TV, EEs from the gastric region responded differently to DMelNV, but in other cases, the specific response was related to cell-subtypes located in the posterior region, such as I-ap-p and II-p.

Next, we evaluated whether generic response genes are up- or down-regulated to perform GO enrichment analysis. Cell-subtype-specific DEGs were also included in this analysis, albeit given the fact that their expression depends on cell-subtype, they were analyzed as a whole (Figure 4). Throughout this work the lists of cell-subtype-specific response genes only included genes that are not present in the generic response. DMelNV and TV exhibited distinct overrepresented terms, indicating that infected cells respond differently to these viruses. However, some similarities between cell infected with DMelNV and TV were noticed. An overrepresentation of terms related to the neuropeptide signaling pathway was found in the TV cell-subtype-specific response genes (Figure 4A), but some neuropeptides such as AstC, CCHa1 and Tk were also differentially expressed in DMelNV infected cells (supplementary file S1). Similarly, translation-related proteins were under-expressed in cells infected with either virus. DMelNV infection led to a wide down-regulation of translation-related genes, mostly ribosomal proteins (Figure 4B and Figure S5A), whereas TV infected cells showed a down-regulation of translation initiation factor eIF3i (Figure S5B).

**Figure 4.**
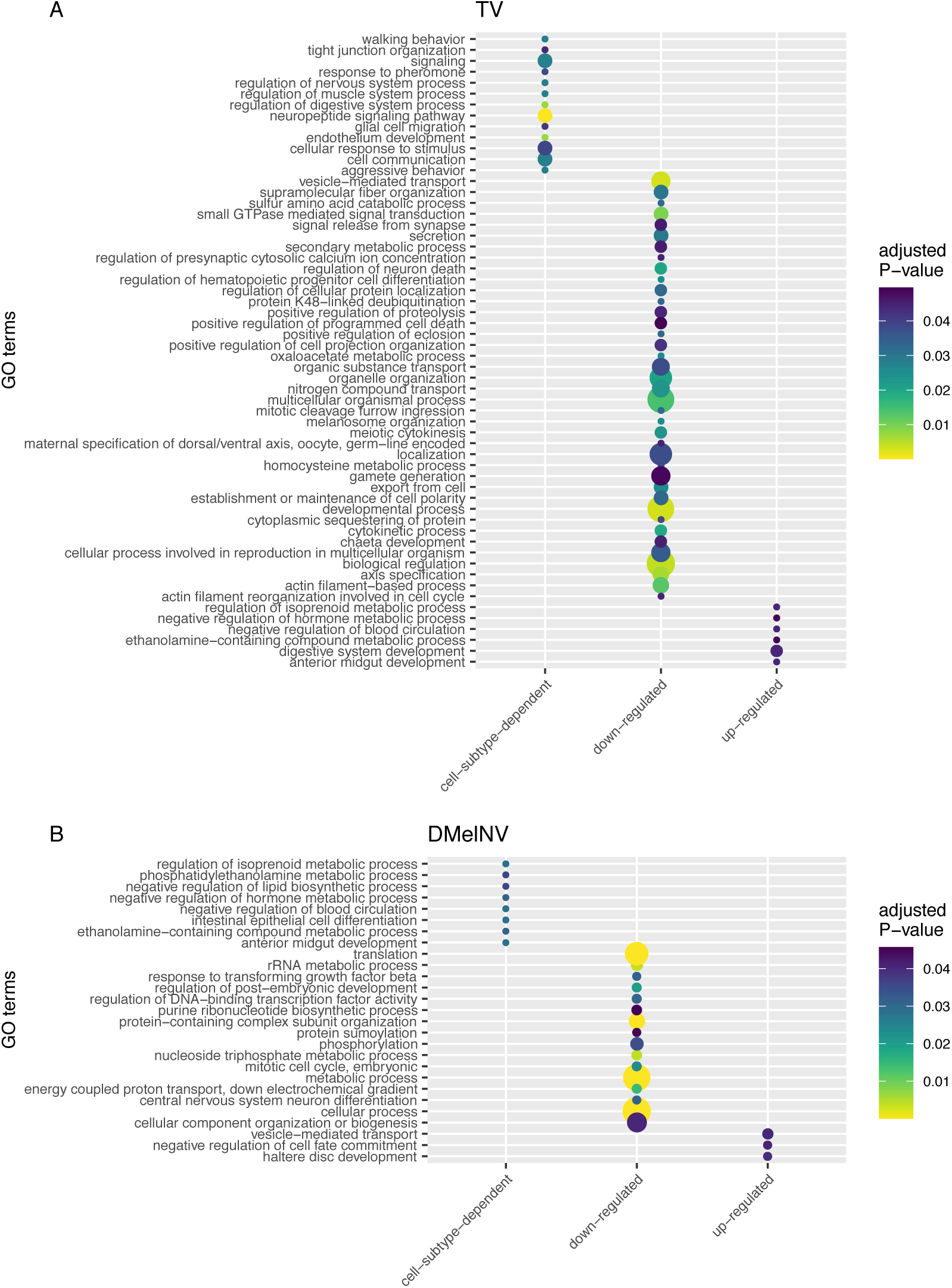
Generic and cell-subtype-specific responses to TV and DMelNV. (A) GO enrichment analysis of DEGs upon infection with TV. The size of the points represents the number of times the respective term was present. (B) GO enrichment analysis of DEGs upon infection with DMelNV for the EEs dataset. The size of the points represents the number of times the respective term was present.

Similar processes were both negatively correlated to DMelNV replication and down-regulated in DMelNV infected cells (Figure 3 and Figure 4B). Although no GO enrichment was found for genes that are positively correlated to DMelNV replication, some genes were present in both positive correlated and up-regulated genes lists, such as Hsc70-3, a chaperone that is also involved in RNAi response (Dorner et al., 2006).

The results obtained for DMelNV were only somewhat reproducible for the midgut atlas dataset, which is not surprising given that new cell types were analyzed. Translation-related processes were enriched in both lists of generic response down-regulated genes and cell-subtype-specific response genes. Viral processes were also overrepresented in cell-subtype-specific responses genes due to the presence of Rack1 in this list. This protein was shown to be necessary for internal ribosome entry site (IRES)-mediated translation of two picorna-like viruses from the family *Dicistroviridae* (Majzoub et al., 2014), and was also shown to interact with the RISC complex (Jannot et al., 2011).

### Robustness of ANOVA tests to determine DEGs

Major limitations of our study include the inability to confidently assign infected cells due to some technical problems, such as the presence of polyadenylated viral RNA in viral particles, and the fact a low number of DMelNV-infected cells were found in the EE dataset. Thus, we sought to determine whether our methodology is robust against these limitations. To this end, random subsets of uninfected cells were assigned as infected and two-ways ANOVA tests were performed as described above. Even at low numbers of cells assigned as infected, a low rate of false-positives was obtained for both generic (infection) and cell-subtype-specific (cell-by-infection interaction term) DEGs (Figure 5), confirming that the two-way ANOVA results are robust against mis-assignments of noninfected cells as infected ones. Notably, the false-positive rate for the midgut atlas dataset was slightly higher and more variable, probably due to the lower number of uninfected cells used to conduct this test in this dataset.

**Figure 5.**
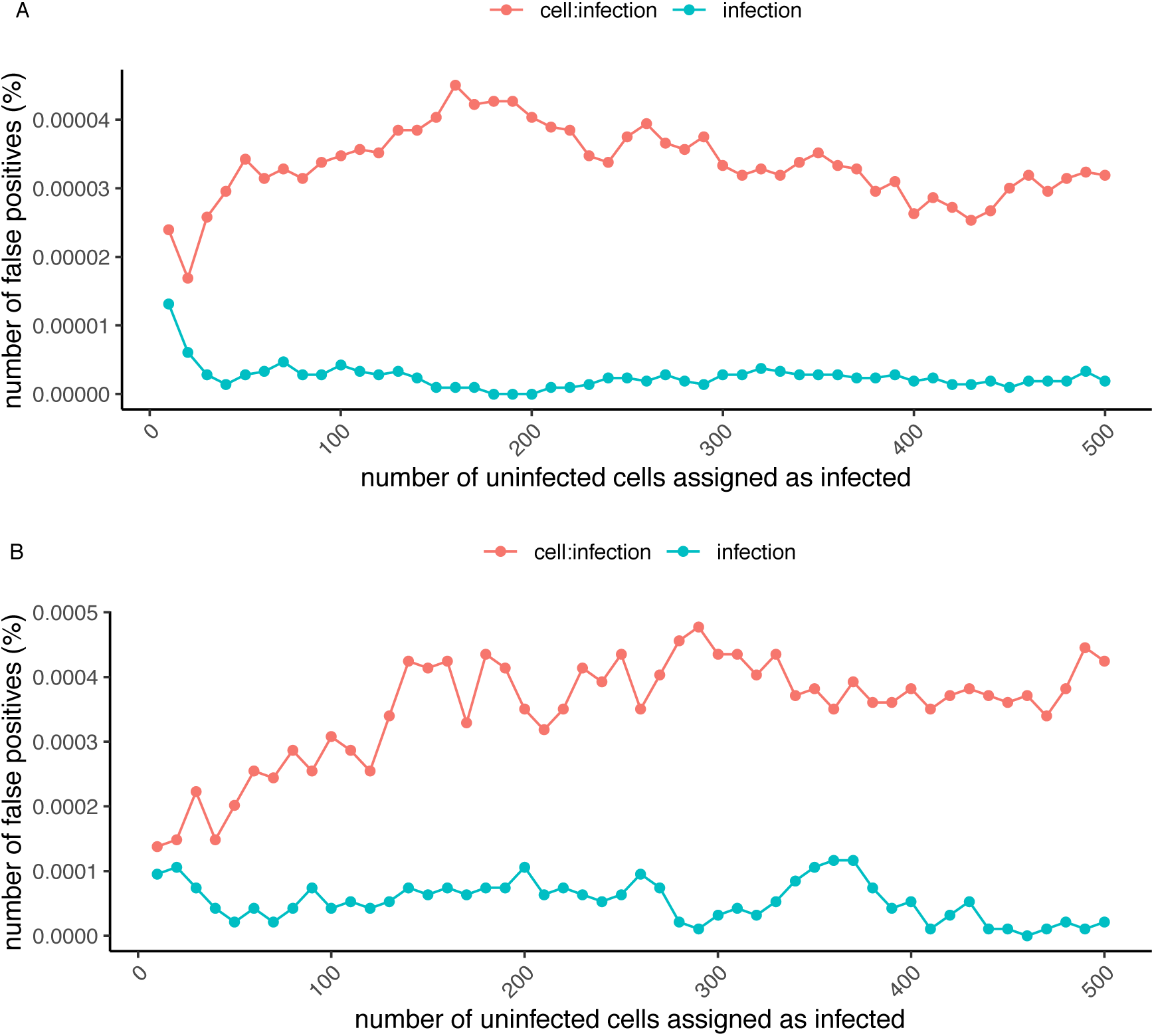
Sensitivity curve for false-positives. (A) Percentage of false-positive DEGs when performing two-way ANOVA tests assigning random uninfected cells as infected for the EE dataset. For each point (coordinates axis), 100 replicates were conducted. (B) Percentage of false-positive DEGs when performing two-way ANOVA tests assigning random uninfected cells as infected for the midgut atlas dataset. For each point (coordinates axis), 100 replicates were performed.

### Mapping generic and cell-subtype-specific responses into *D. melanogaster* interactome

A network biology approach was taken to study the effects of the DEGs on the host transcriptome. In this approach, the *D. melanogaster* interactome is represented by a network where genes (nodes) are connected to each other by edges to represent their interactions. Topological parameters of a predicted *D. melanogaster* interactome network were computed to investigate whether DEGs constitute essential nodes in the network. Two parameters were estimated: degree, the number of interactions of a gene; and betweenness, which is approximately the number of shortest paths that cross a gene in the network. Essential genes very likely are defined by either a high degree (hubs) or high betweenness (bottlenecks) (Cho et al., 2012; Ahmed et al., 2018). The degree of DEGs were analyzed by comparing their degree distribution with that of the complete fly interactome. Figure 6 and Figure S6A shows the power-law fit to the log-degree distribution of the complete interactome network and subnetworks constructed using only DEGs found in the EE and midgut atlas dataset, respectively. The values of the critical exponent, mean degree and mean betweenness for each network are shown in Table 1. The power-law critical exponent estimated for the DEGs was significantly smaller than for the whole interactome only for DMelNV down-regulated genes in the EE dataset (*P* = 0.017) and DMelNV negatively correlated genes in the midgut atlas dataset (*P* = 1.77×10^−3^). These results indicate that DMelNV infection led to an under-expression of hub genes with a high number of interactions. Next, the betweenness of DEGs were compared to that from the complete interactome. In this case, genes that were under-expressed upon infection with either DMelNV (in both datasets) and TV showed a significant higher betweenness than expected from the whole genes of the fly interactome. This indicates that while DMelNV infection leads to a down-regulation of both hub and bottleneck genes, TV infection leads only to a down-regulation of bottleneck genes.

**Table 1.**
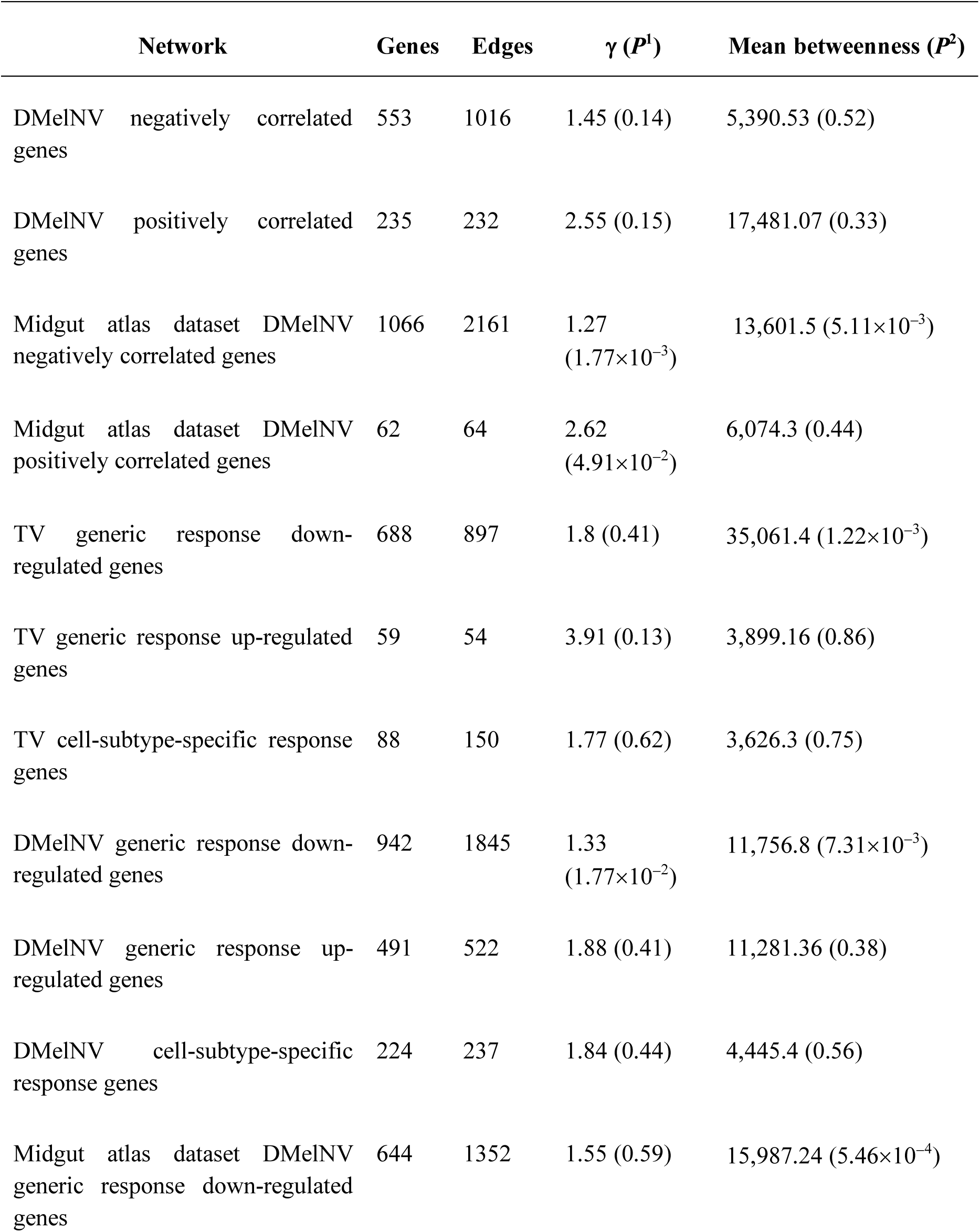
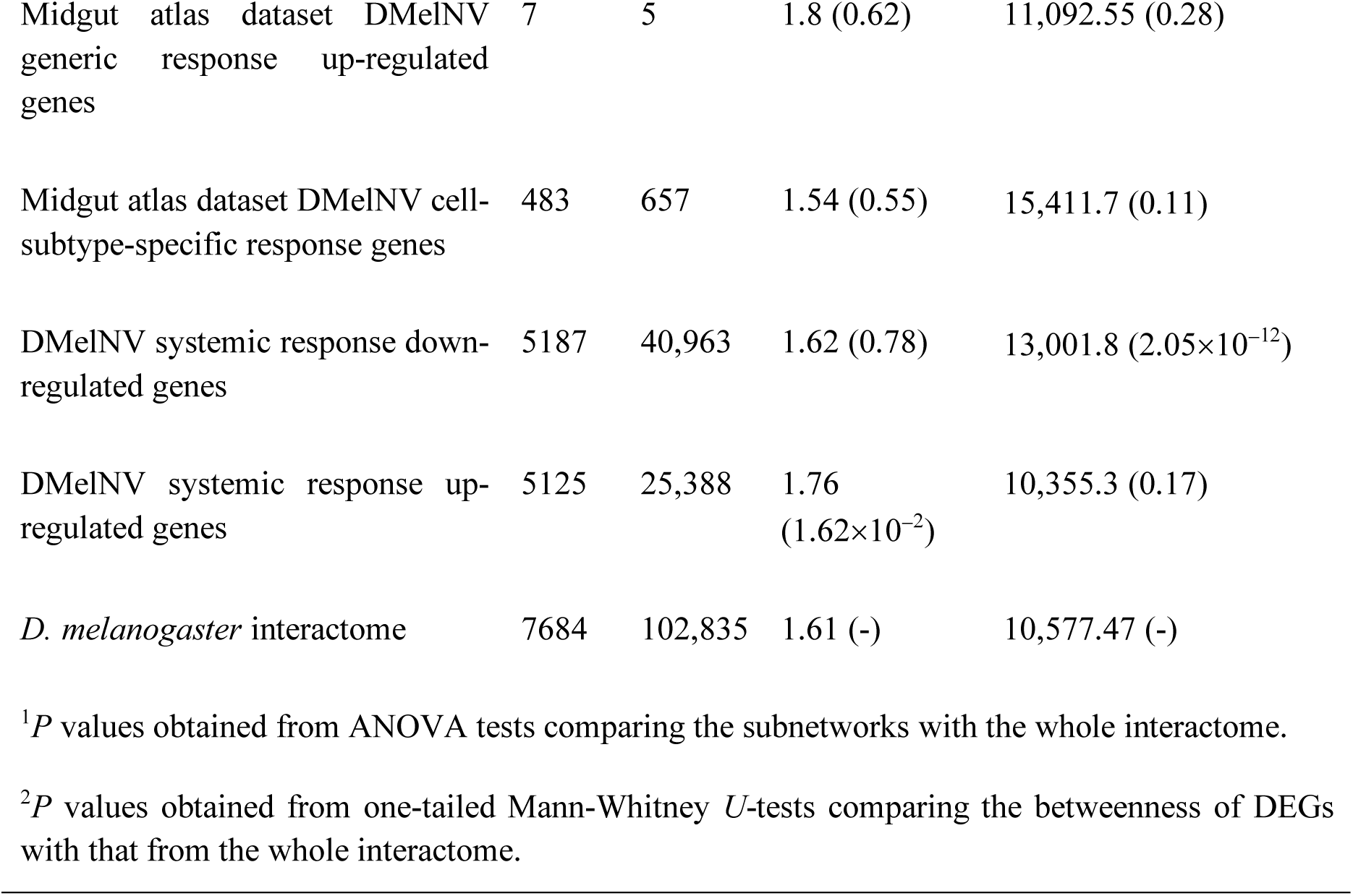
Topological network parameters estimated for the *D. melanogaster* network and DEGs subnetworks.

**Figure 6.**
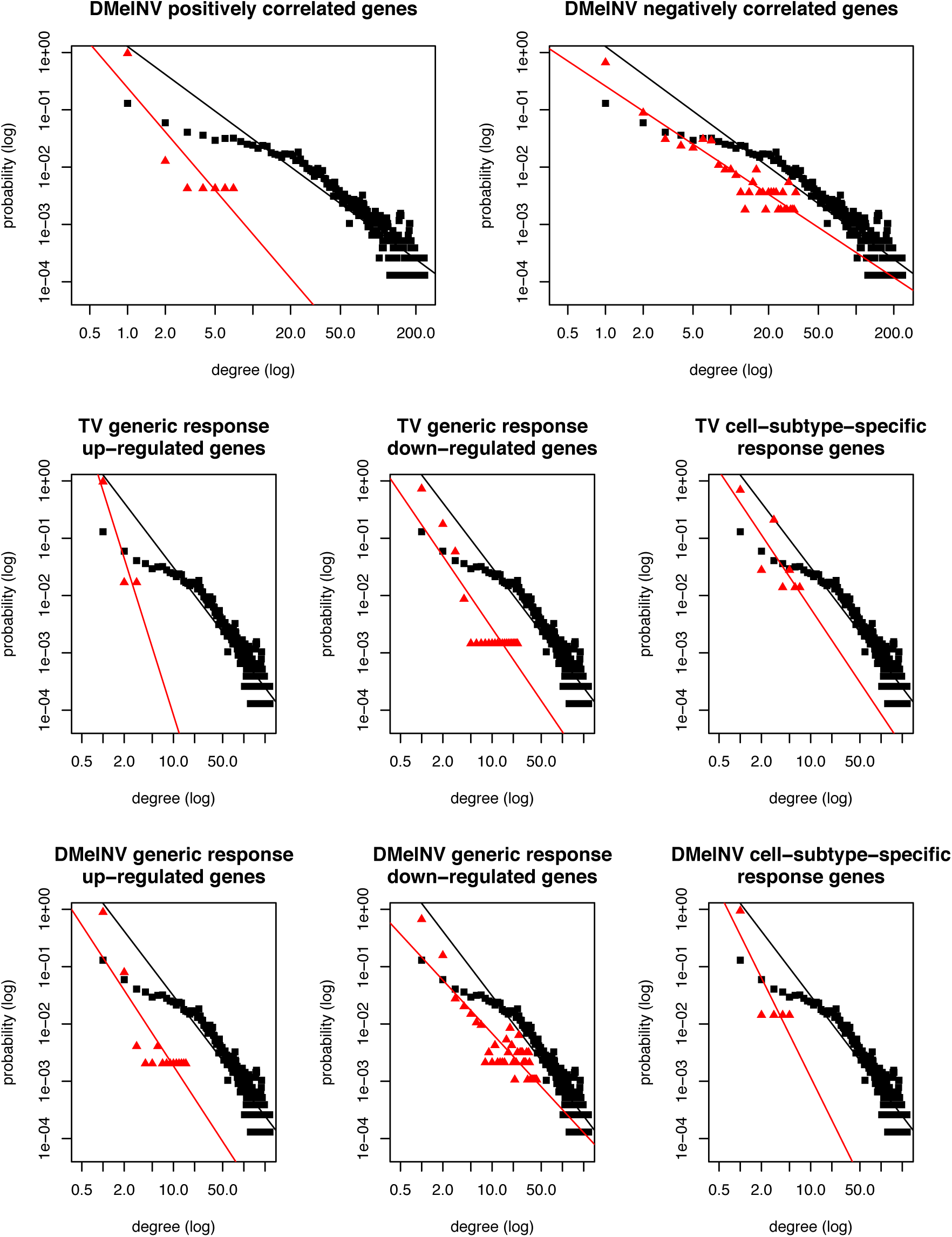
Mapping generic and cell-subtype-specific responses into *D. melanogaster* interactome. Log-degree distributions of the whole *D. melanogaster* interactome (black) and subnetworks constructed with DEGs (red) for cells infected with TV or DMelNV. Regression lines represent the critical exponent of the power-law.

### The cellular response to DMelNV, but not the systemic response, is characterized by a down-regulation of hub genes in the *D. melanogaster* interactome

So far, only the cellular response to viral infection was evaluated here. To better understand the response to viral infections in a systemic manner, we analyzed a publicly available time series bulk RNA-seq data from DMelNV-infected *D. melanogaster* (Lopez et al., 2018) and compared it with our previously obtained results. It is important to note, however, that the cellular response was obtained by comparing infected/uninfected cells but we are unaware whether uninfected cells were exposed to viruses. Another limitation of this approach is that bulk RNA-seq of infected flies will average the gene expression for all cell types, including infected and uninfected cells. Five and four genes were simultaneously down- and up-regulated, respectively, in both cellular and systemic responses (supplementary file S1). Surprisingly, two genes, Hsc70-3 and Hsc70Cb, were under- and over-expressed in the systemic and cellular response, respectively, albeit in the systemic response they were only under-expressed at 30 days post-eclosion. As mentioned above, Hsc70-3 was also correlated to the replication level of DMelNV. Ribosomal proteins, however, were widely down-regulated in the cellular response, but a few of them were up-regulated in the systemic response. A similar pattern of down- and up-regulation in the cellular and systemic response, respectively, was noticed for other proteins such and NPF and Tk (supplementary file S1). The degree and betweenness of the systemic response DEGs were calculated. In contrast to the cellular response, we only found a significant modulation of bottleneck genes for the systemic response to DMelNV (Figure S6B and Table 1), suggesting that interference with hub genes in the fly interactome is exclusive to the cellular response.

## Discussion

By analyzing scRNA-seq data from *D. melanogaster* EEs, we showed that two picorna-like viruses, DMelNV and TV, employ different strategies regarding replication and alteration of host transcription. DMelNV was also found on a variety of cell types in a second scRNA-seq data of the whole fly midgut, and possibly, both DMelNV and TV infect other cell types not included in this study. Whereas DMelNV is known to be a non-pathological enteric virus, the biology of TV remains unknown as this virus has been recently described by a metagenomic approach (Webster et al., 2015). Here, we showed that TV capacity to infect and its replication level on EEs depended on the particular cell-subtype being infected, and also, its accumulation was generally low compared to DMelNV (Figure 2). Up to 60% of the I-p^AstA^ subtype, located in the posterior region of the gastrointestinal tract, was infected with TV, while a low percentage of EEs from the gastric region was infected with this virus. TV accumulation was also significantly higher in the I-p^AstA^ and II-p subtypes while significantly lower in the II-a, III and II-m subtypes. On the other hand, DMelNV replication level on EEs was either low or high independent on cell-subtype, where the percentage of viral RNA in the cells sometimes exceeded 50% and was up to ∼80% (Figure 2). Although its prevalence varied from cell subtype in the midgut atlas dataset, we failed to detect any significant difference in its log-expression values across these subtypes, suggesting that the observed difference in percentage of viral RNA and accumulation (Figure S2) may be due to other factors such as sample size. Similar to DMelNV, a wide range in viral load/transcripts in infected cells was observed for other RNA viruses such as vesicular stomatitis virus (VSV) (Zhu et al., 2009), dengue and zika viruses (Zanini et al., 2018), poliovirus (Schulte and Andino, 2014), foot-and-mouth disease virus (Xin et al., 2018), and influenza A virus (IAV) (Heldt et al., 2015; Steuerman et al., 2018; Russell et al., 2018).

Cells infected with DMelNV, independent on subtype, showed an under-expression of various proteins related to translation, mostly ribosomal proteins (Figure 4B and Figure S3). This indicates that DMelNV infected cells undergo a widespread shutoff of the translational machinery, possibly to tamper its replication. Alternatively, DMelNV targets only specific ribosomal genes as an attempt to suppress the translation of host RNA while enhancing the translation of its own proteins, and therefore, enhancing its own replication. The fact that disruption of essential genes in the interactome might diminish the infection process of pathogens, in accordance with the centrality-lethality rule (Jeong et al., 2001; Ahmed et al., 2018), corroborates with the first hypothesis; whereas the fact that some cells exhibited a large amount of DMelNV RNA and the expression of some ribosomal protein genes is negatively correlated to DMelNV replication level corroborates with the latter hypothesis. Interestingly, previous micro-array and bulk RNA-seq studies were unable to find a similar under-expression of ribosomal genes in DMelNV-infected flies (Cordes et al., 2013; Lopez et al., 2018). In fact, some ribosomal proteins were over-expressed in the bulk RNA-seq data (supplementary file S1). In addition, genes that were under-expressed in DMelNV-infected cells likely constitute essential hubs and bottlenecks in the fly’s interactome (Figure 6 and Table 1). On the contrary, analysis of the bulk RNA-seq data showed only a significant down-regulation of bottleneck genes (Table 1). These differences highlight the distinction between cellular and systemic response to viral infections.

This question has been recently addressed in details in a scRNA-seq study using mice infected with IAV (Steuerman et al., 2018). It was noted an over-expression of a type I interferon (IFN) gene module when comparing uninfected cells from an IAV infected mouse (called bystander cells) with cells from a virus-free mouse, whereas the over-expression of this module was less noticeable when comparing IAV-infected cells with bystander cells (Steuerman et al., 2018). Likewise, it was shown that both severe acute respiratory syndrome coronavirus-2 (SARS-CoV-2)-infected cells and bystander cells also exhibit an up-regulation of interferon-stimulated genes (Ravindra et al., 2020). Similar to DMelNV, several plant viruses were found to interact with hubs and bottleneck genes of the *Arabidopsis thaliana* interactome (Rodrigo et al., 2012). These hubs are mostly related to the plant immune response, which indicates that a substantial reprogramming of the host transcriptome is necessary to initiate an immune response. In our case, we failed to detect any considerable up-regulation of immune genes in the cellular response to either TV or DMelNV. Interestingly, the time course analysis of the response to DMelNV found that expression of immune related genes increased overtime (Lopez et al., 2018). As the time of infection is an important factor to regulation of the host transcriptome, and the scRNA-seq data used here is from 5-7 days old flies, we may have failed to detect a more substantial change to the host transcriptome. The up-regulation of immune genes in the bystander response to IAV (Steuerman et al., 2018) and SARS-CoV-2 (Ravindra et al., 2020) and in the bulk RNA-seq analysis of DMelNV-infected flies (Lopez et al., 2018) indicates that activation of immunology-related genes is triggered as a systemic response to viral infections.

In contrast, we failed to detect any perturbation of hub genes in the interactome for TV infection. One reason why modulation of hub genes upon TV infection was not detected may be that we only analyzed the cellular response to TV infection, and such alteration of hub elements of the host transcriptome may still happen in a systemic manner, and as such, may only be detected by comparing virus-free flies with infected ones. Again, it is worth noticing that in our case, data was generated from five-seven days old flies, and the time course of infection may be crucial to the modulation of host transcriptome. However, TV infection led to a down-regulation of bottleneck genes. Despite the fact that ribosomal genes were not under-expressed in TV-infected cells, we found a down-regulation of eIF3i and a surprisingly up-regulation of two mitochondrial ribosomal genes. Many viruses from the order *Picornavirales* are known to target initiation factor proteins, usually through cleavage of the protein, to shutoff the host translational machinery and increase its protein synthesis (Garrey et al., 2010; Sommergruber et al., 1994; Gradi et al., 2004).

We also investigated the cell-subtype-specific response to DMelNV and TV infection. No evidence for modulation of hub or bottleneck genes was found for cell-subtype-specific response DEGs. This suggests that in order to infect additional cell-subtypes, viruses need to alter/exploit auxiliary pathways. Interestingly, some neuropeptide hormones such as AstC, CCHa1, CCHa2, and Tk are differentially expressed in cells infected by DMelNV or TV in a cell-subtype-specific manner. In fact, the neuropeptide signaling pathway process was overrepresented in cell-subtype-specific DEGs upon TV infection.

Last, given the presence of unacknowledged viruses in public scRNA-seq data, we hypothesize that viruses may be confounding factor in these kinds of experiments. In the data analyzed here, viruses did not seem to cluster based on infection status (Figure 1 and Figure S2A). However, we have no information of which uninfected cells are responding to virus infections as bystanders, adding more hidden confounding factor to these analyses. One possibility to mitigate this problem would be to remove from the analysis all immune-related genes or genes which are correlated to viral load. A similar approach, where genes correlated to the top two PCs composed by anti-viral and inflammatory genes were omitted from the analysis, was performed to cluster pulmonary cells from IAV-infected and uninfected mice without the possible confounding effects of anti-viral genes (Steuerman et al., 2018).

Here, through analysis of *in vivo* scRNA-seq data, we show the similarities and differences in the replication and infection strategies of two *D. melanogaster* viruses. This powerful technique can be used to monitor both pathogen(s) and host transcriptomes simultaneously, analogous to dual RNA-seq strategies. We also highlight differences between the cellular and systemic responses to DMelNV infection. The results obtained here should provide models that could be useful for other invertebrate viruses.

## Acknowledgements

This study was financed in part by the Coordenação de Aperfeiçoamento de Pessoal de Nível Superior - Brasil (Capes) - Finance Code 001, and by Generalitat Valenciana grant PROMETEU2019/012, CSIC grant PIE202020E94 and Spain Agencia Estatal de Investigación - FEDER grant PID2019-103998GB-I00 to S.F.E. T.N. and F.L.M received a PQ-fellowship from CNPq.

## Author Contributions

J.M.F.S. performed the analyses and wrote the draft. T.N., F.L.M. and S.F.E supervised the study. All authors revised the manuscript.

---

**Figure S1.**
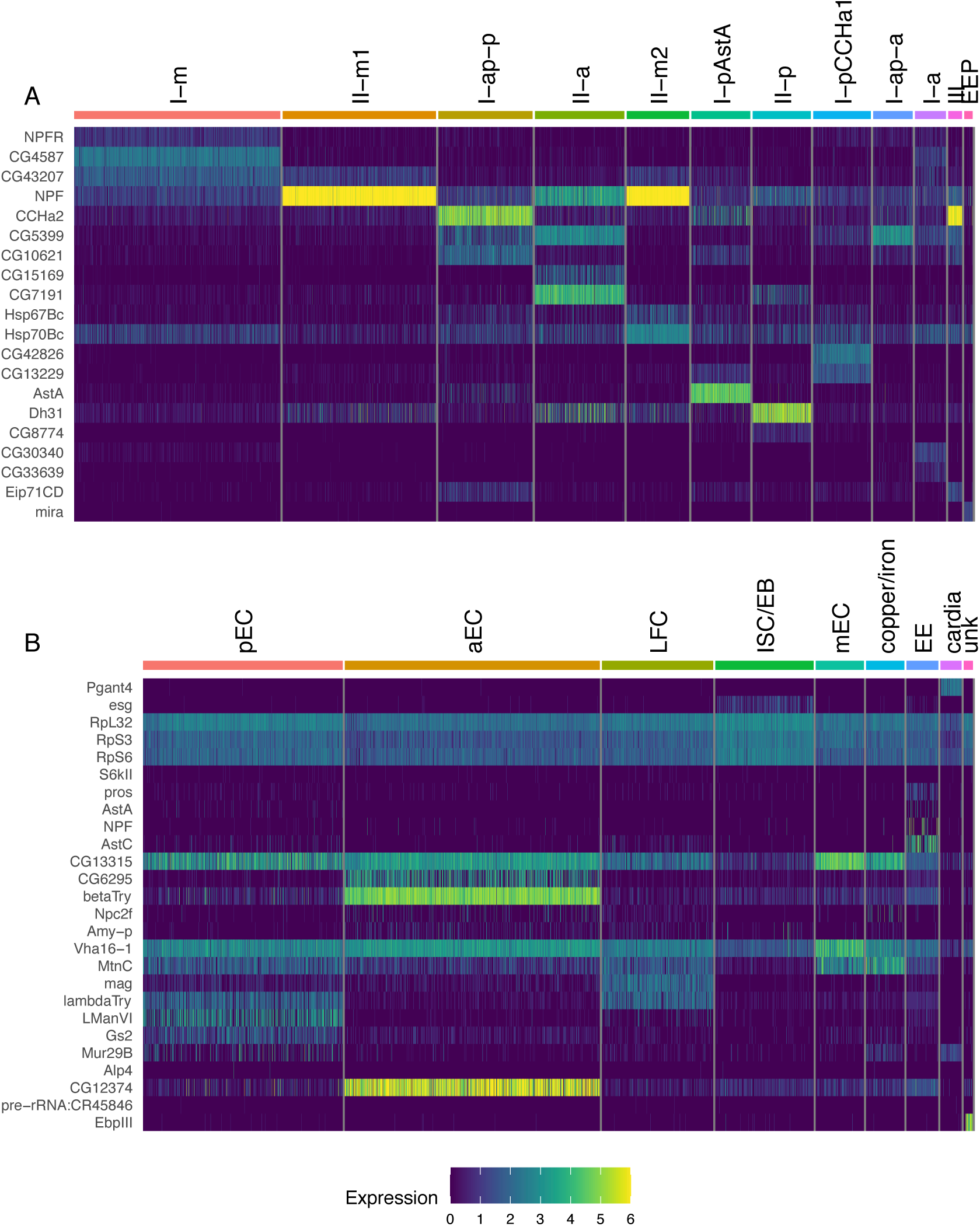
Marker genes used to identify clusters. (A) Heatmap of marker genes for the EEs dataset. (B) Heatmap of marker genes for the midgut atlas dataset.

**Figure S2.**
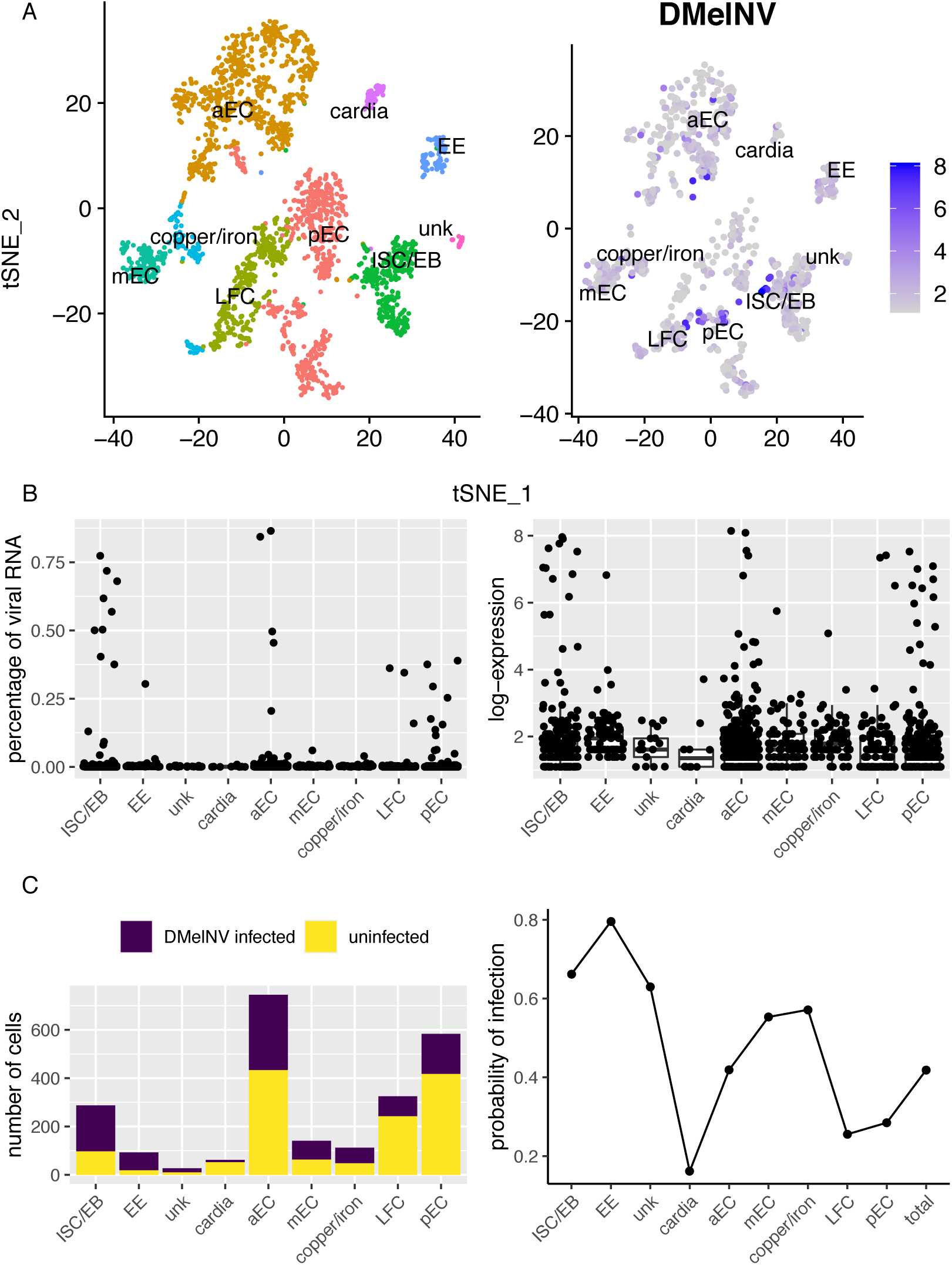
Detection of DMelNV on a *D. melanogaster* midgut scRNA-seq dataset. (A) t-SNE reduction plots with identified clusters (left); and cells infected with DMelNV along with their log-expression level (right). (B) Boxplots showing the percentage of viral RNA (left) and log-expression values (right) of DMelNV for each midgut cell major subtype. Each point represents an individual cell. (C) Number (left) and percentage (right) of cells from each subtype infected with DMelNV.

**Figure S3.**
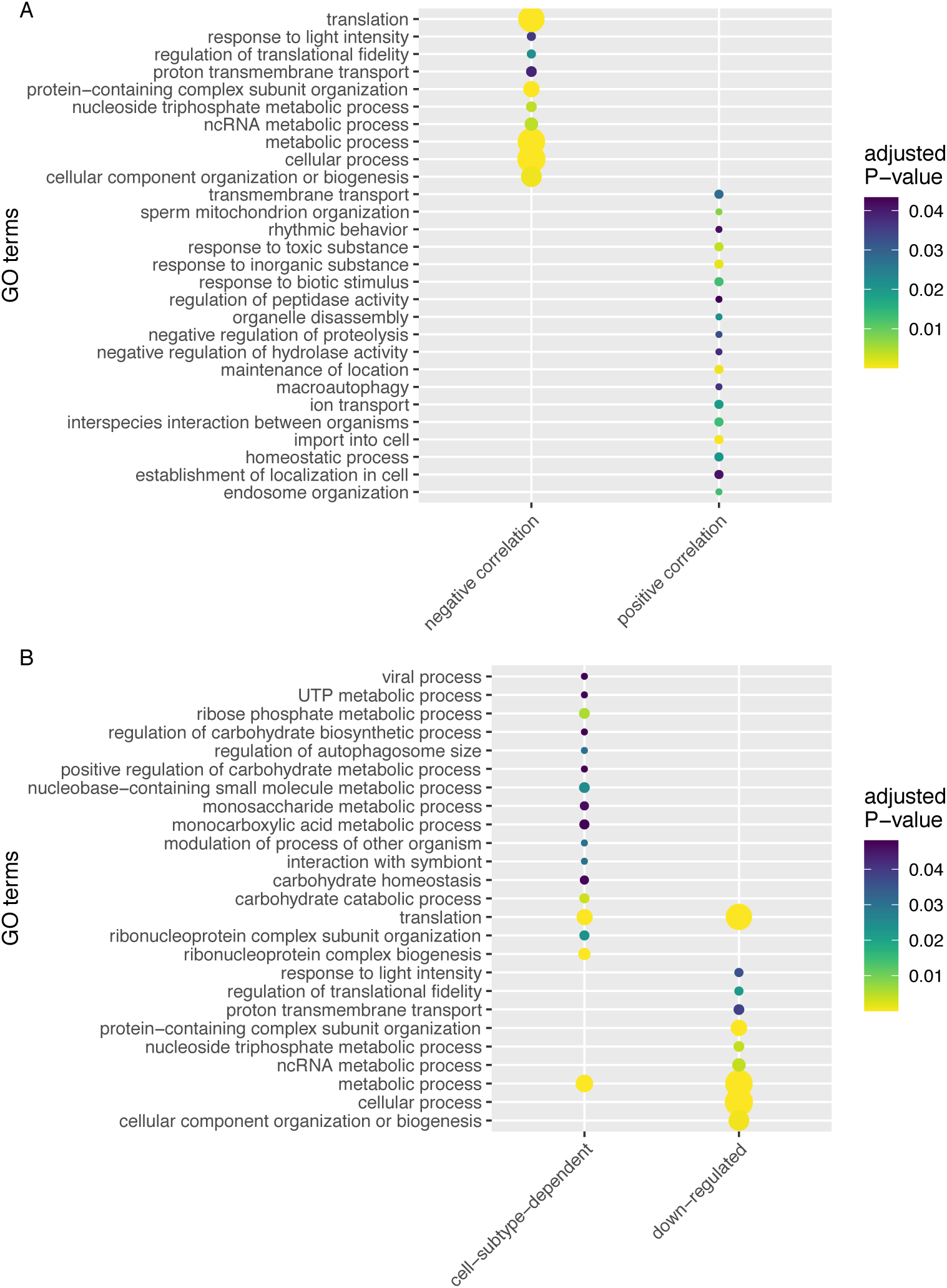
DMelNV replication correlation to gene expression and DEGs for the midgut atlas dataset. (A) GO enrichment analysis of genes which expression is correlated to the replication level of DMelNV for the midgut atlas dataset. The size of the points represents the number of times the respective term was present. (B) GO enrichment analysis of DEGs upon DMelNV infection for the midgut atlas dataset. The size of the points represents the number of times the respective term was present.

**Figure S4.**
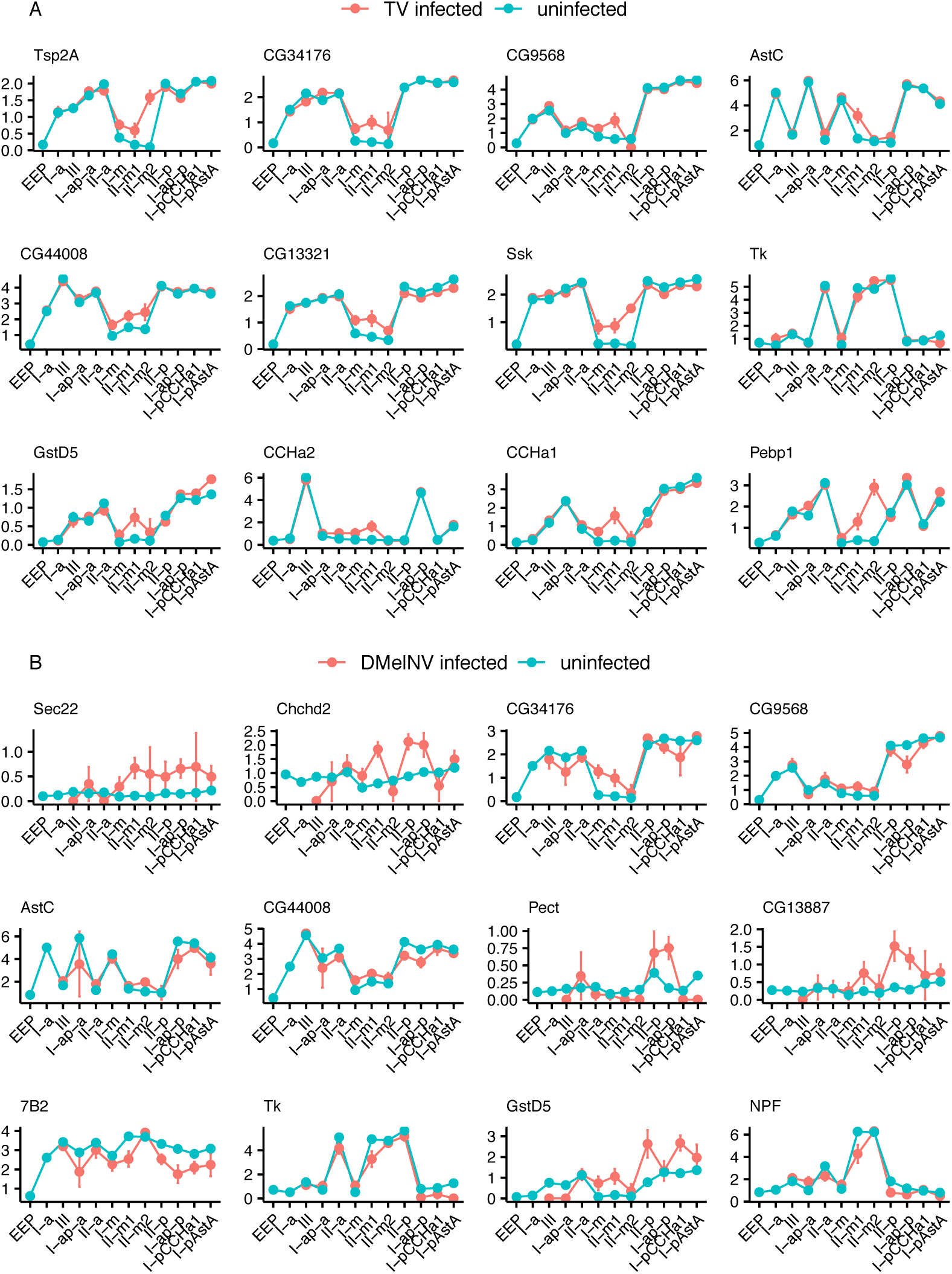
Cell-subtype-specific response to TV and DMelNV. (A) Cell-subtype-specific response genes to TV infection. Twelve DEGs with the lowest adjusted *P* < 0.05 for the interaction between cell-subtype and infection status in the ANOVA analysis are shown. (B) Cell-subtype-specific response genes to DMelNV infection. Twelve DEGs with the lowest adjusted *P* < 0.05 for the interaction between cell-subtype and infection status in the ANOVA analysis are shown.

**Figure S5.**
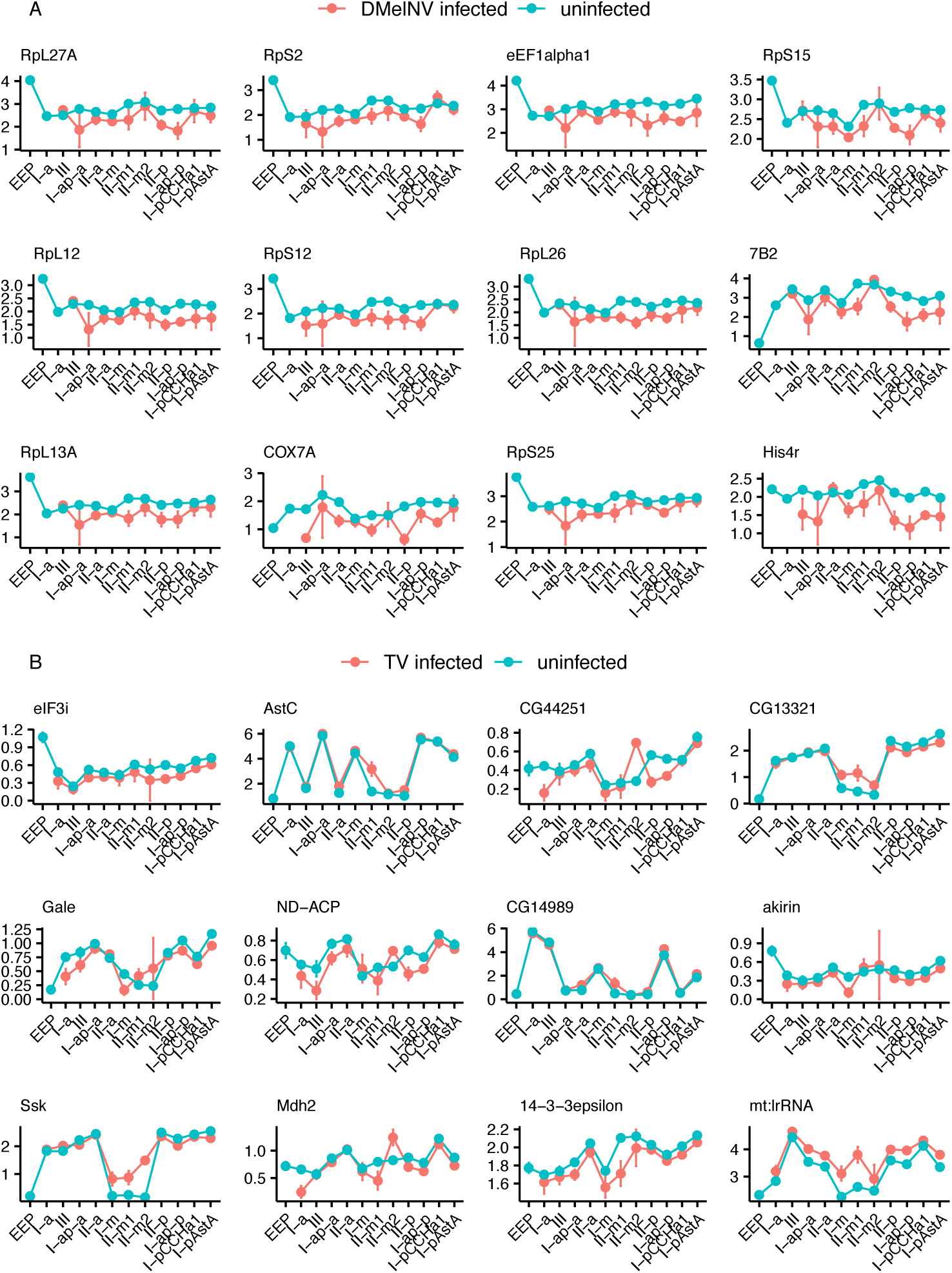
Generic cellular response to DMelNV and TV. (A) Generic cellular response genes to DMelNV infection. Twelve DEGs with the lowest adjusted *P* < 0.05 for the interaction between cell-subtype and infection status in the ANOVA analysis are shown. (B) Generic cellular response genes to TV infection. Twelve DEGs with the lowest adjusted *P* < 0.05 for the interaction between cell-subtype and infection status in the ANOVA analysis are shown.

**Figure S6.**
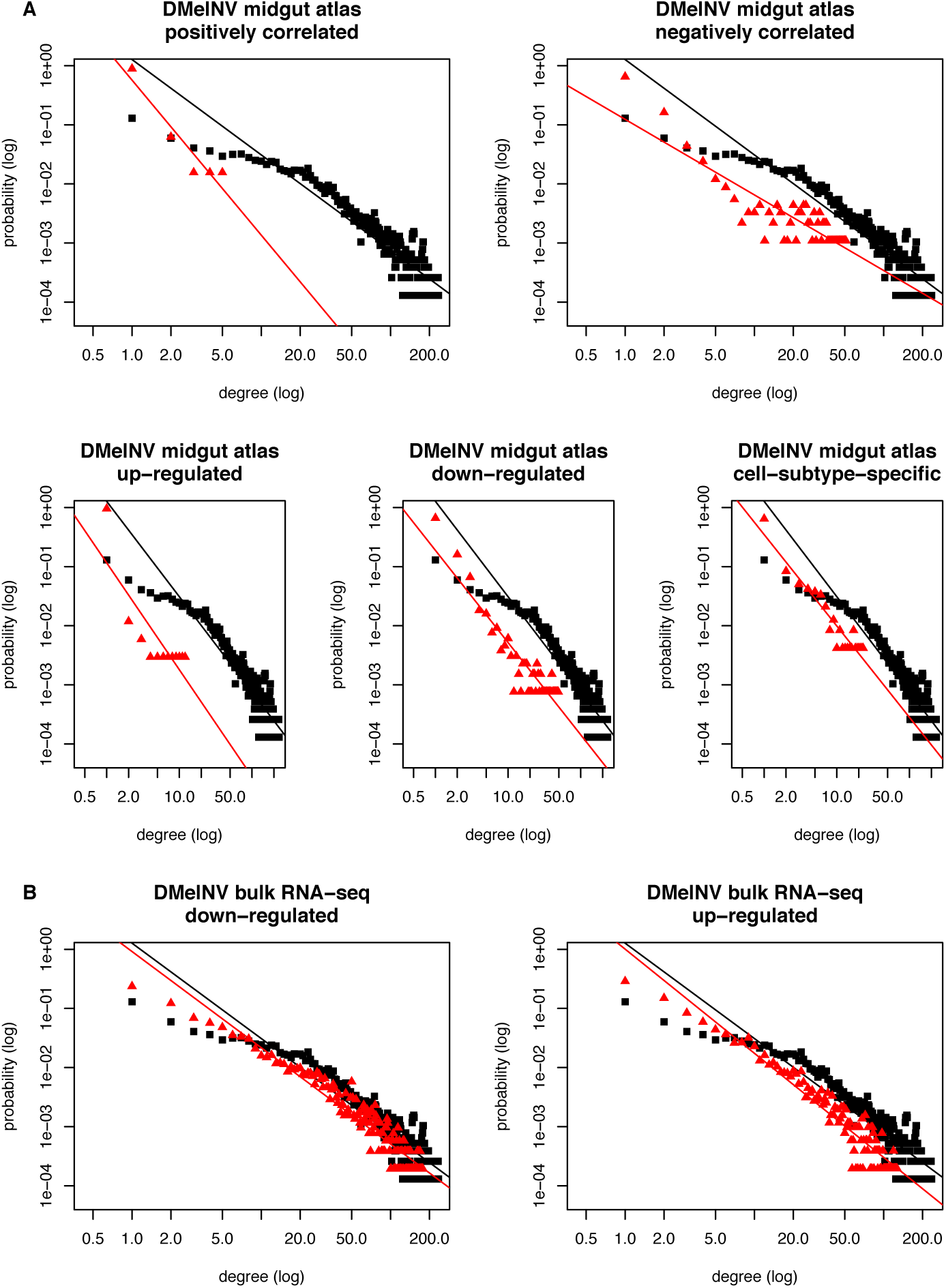
Mapping of DEGs found in the midgut atlas and bulk RNA-seq datasets into *D. melanogaster* interactome. (A) Log-degree distributions of the whole *D. melanogaster* interactome (black) and subnetworks constructed with DEGs (red) detected in the midgut atlas dataset. The slopes of the regression lines correspond to the critical exponent of the power-law. (B) Log-degree distributions of the whole *D. melanogaster* interactome (black) and subnetworks constructed with DEGs (red) detected in the bulk RNA-seq analysis of DMelNV infected flies (Lopez et al., 2018). The slopes of the regression lines correspond to the critical exponent of the power-law.

## STAR Methods

### Data collection

Raw sequencing data was downloaded from NCBI SRA database (https://www.ncbi.nlm.nih.gov/sra; BioProject accession: PRJNA547484). Briefly, this data was generated by following 10× Genomics GemCode protocol (Zheng et al., 2017) using ∼8000 EEs harvested from the midgut of ∼200 female fruit flies (CG32547-GAL4 > GFP line) aged between five and seven days (Guo et al., 2019). GFP was used to sort cells with a FACS Aria III sorter (BD Biosciences) and sequencing was performed at an Illumina X10 platform. Additionally, a fruit fly scRNA-seq dataset from the entire midgut was also obtained through SRA (BioProject accession: PRJNA493298). Only data generated by 10× technology was analyzed. Briefly, guts from seven days old female flies (esg-sfGFP/+, pros-GAL4 > RFP/+ line) were dissected and libraries were prepared following 10× Genomics GemCode protocol (Hung et al., 2020). Two technical replicates for two samples were prepared for this data, resulting in a total of four libraries.

### Filtering and cluster generation

BLASTn (Altschul et al., 1990) searches of the reads against the virus RefSeq database were conducted as a preliminary analysis to identify possible viruses in this sample. Three picorna-like viruses were found: DCV (accession AF014388), DMelNV (accession GQ257737) and TV (accession KP714072). Barcode processing and gene counts were performed with cellranger v3.1.0 (10× Genomics, CA, USA) using an edited *D. melanogaster* reference genome (accession: GCA_000001215.4) that included the viral sequences. Downstream analysis was performed with Seurat v3.1.4 (Stuart et al., 2019). 4994 cells containing between 200 and 3000 detected genes and < 5% mitochondrial genes were retained. Scaling and normalization were performed with the SCTransform function (Hafemeister and Satija, 2019), with mitochondrial genes and virus counts regressed out. Principal component analysis (PCA) and *t*-distributed stochastic neighbor embedding (t-SNE) reductions were performed with top 20 principal components (PCs) and clusters were generated at a 0.4 resolution. Clusters were identified using marker genes (Guo et al., 2019) as shown in Figure S1A.

The same analysis was performed for the midgut atlas dataset, albeit with some minor differences. Most importantly, in this dataset only DMelNV could be identified. Cellranger v3.1.0 was run separately for each library. Technical replicates counts were merged, and the two samples were integrated using SCTransform normalized counts with FindIntegrationAnchors and IntegrateData functions in Seurat v3.1.4. After filtering, 2375 cells containing between 200 and 3000 detected genes and < 25% mitochondrial genes remained. All integration steps, PCA and t-SNE reduction were performed with top 50 PCs and clusters were generated at a 1.0 resolution. Clusters were identified with a set of marker genes (Hung et al., 2020) as shown in Figure S1B.

### Virus infection/replication analyses

Only cells with at least 2 normalized viral UMIs were considered infected. This approach was taken since cells with low amount of viral RNA may represent false-positives caused by contamination of viral particles, contamination of RNA that leaked from other cells during lysis or other technical problems. By using this criterion, we effectively set the minimum frequency of viral UMIs to 3.1×10^−4^ and 2.8×10^−4^ for DMelNV and TV in the EE dataset, respectively, and to 3.9×10^−4^ for DMelNV in the midgut atlas dataset. These thresholds are on the same ball park than those used in other studies with IAV (2×10^−4^ - 1.4×10^−3^) (Steuerman et al., 2018; Russell et al., 2018). The percentage of viral RNA was calculated using raw UMI counts and log-expression values were obtained using normalized values. One-way ANOVA tests were performed to investigate whether the accumulation of DMelNV and TV are influenced by cell subtype using R v3.6.2. Significant results were further investigated by performing Tukey’s HSD post *hoc tests* using the agricolae R package v1.3-2 (https://CRAN.R-project.org/package=agricolae) in R v3.6.2.

### Gene expression analyses

All gene expression analyses were performed on a list of 1460 genes that are expressed in at least 10% of the cells in each cluster. For the midgut atlas dataset, a list composed by 307 genes was generated. Partial correlations controlling for cell subtype were performed for each virus/gene pair using the ppcor R package v1.1 (Kim, 2015). For both DMelNV and TV, two-way ANOVA tests were conducted on the expression level of each gene on the list accounting for cell subtype, infection status and the interaction of these two orthogonal factors. The average log-fold change of the gene expression across all clusters was used to determine whether a gene was up or down regulated. For both partial correlation and two-way ANOVA tests, only genes with a Bonferroni-corrected *P* < 0.05 were analyzed. GO enrichment analyses for biological process were performed with BiNGO v3.0.4 (Maere et al., 2005) plugin for Cytoscape v3.7.2 (Shannon et al., 2003) using the GO-basic and gene annotation files from FlyBase (https://flybase.org/; FB2020_02 release). To simplify the output, child terms were removed when the parent was also present, with the exception of translation (GO:0006412) and neuropeptide signaling pathway (GO:0007218). All DEGs are available at supplementary file S1.

### Robustness of DEGs analysis

To investigate the possible effect of uninfected cells being erroneously assigned as infected, we conducted a step-wise analysis of the false-positive rate by randomly assigning uninfected cells as infected and performing two-way ANOVA tests as described above. For each step, 100 two-way ANOVA tests were carried out with 10 to 500 cells randomly assigned as infected, using an incremental step of 10. Note that the tests were executed on the lists of 1460 and 307 genes for the EE and midgut atlas datasets, respectively, meaning that for each step, 146,000 and 30,700 two-way ANOVA tests were performed for each dataset.

### Gene network analyses

A high quality predicted interactome of *D. melanogaster* was downloaded from http://drosophila.biomedtzc.cn v2018_01 (Ding et al., 2020). Gene interaction networks were analyzed as undirected graphs with the igraph R package v1.2.5 (Csardi and Nepusz, 2006). The degree probability distribution and the betweenness of each node was computed for the interactome network. Then, a linear regression on the degree probability distribution in the log-log space was computed to obtain the critical exponent, γ, of the power-law fit. For each list of DEGs, the corresponding subnetwork was extracted from the complete interactome and its critical exponent was obtained as described above. Next, we performed *t*-tests between the critical exponent of the full interactome and each subnetwork to test for significant differences in the degree distribution between the subnetwork and complete interactome. The betweenness of DEGs were computed using the complete network. One-tailed Mann-Whitney *U*-tests were then performed to compare the betweenness of the DEGs with that obtained from all genes in the network, considering the upper tail of the distribution.

### DMelNV infection bulk RNA-seq data analysis

A list of DEGs from DMelNV-infected female flies was obtained from Lopez et al. (2018). This data contains DEGs detected at two, 10, 20, and 30 days post-eclosion. Briefly, this data was generated by stablishing DMelNV-infected white-eyed flies (*w*^*1118*^; Vienna Drosophila Resource Center, Vienna, Austria) stocks via fecal-oral infection. RNA extraction was performed with TRIzol reagent (ThermoFisher Scientific, Waltham, MA) on triplicates for each time point and samples were sequenced at an Illumina HiSeq system platform. FPKM values were used to determine DEGs. Here, DEGs from each time point were pooled together and gene network analysis was performed for up- and down-regulated genes as described above.

### Resource availability

All scripts used in this work are available at https://github.com/jmfagundes/dmelscrna.

